## Supplemental Information for "Machine Learning Differentiates Enzymatic and Non-Enzymatic Metals in Proteins"

### Supplemental Figures

**Supplemental Figure S1: Enzyme types for enzymatic sites in the dataset.** (A) and test-set (B). Collected from E.C. number or PDB Classification. Marked as 'other' when neither could be used to identify enzyme type.

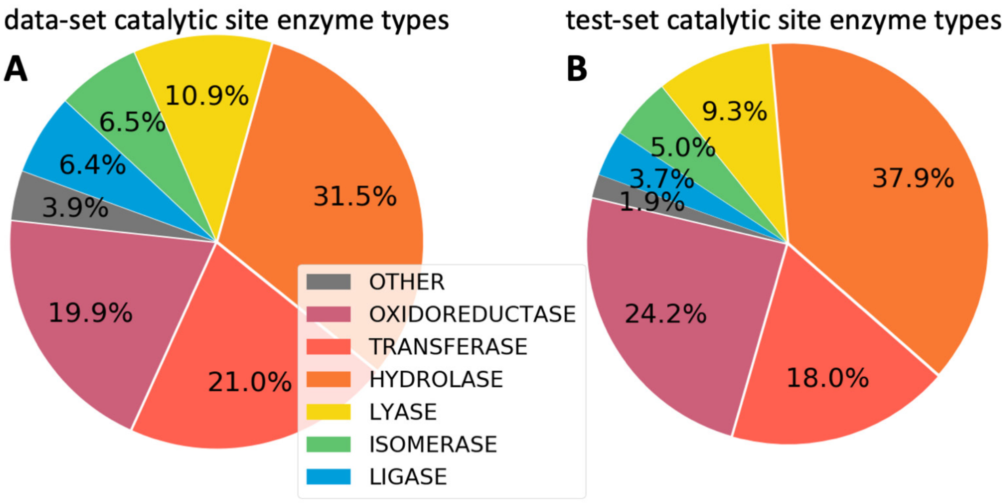


**Supplemental Figure S2: Sphere and Sum features.** 2 dimensional representation of sphere (A) and shell (B) regions. Different wedges represent regions where any atom with an α-carbon within that region will lead to that residue being used for the calculation. Orange wedge represents 3.5 Å cutoff, Yellow wedge represents 5.0 Å cutoff, green wedge represents 7.5 Å cutoff, and blue wedge represents 9 Å cutoff.

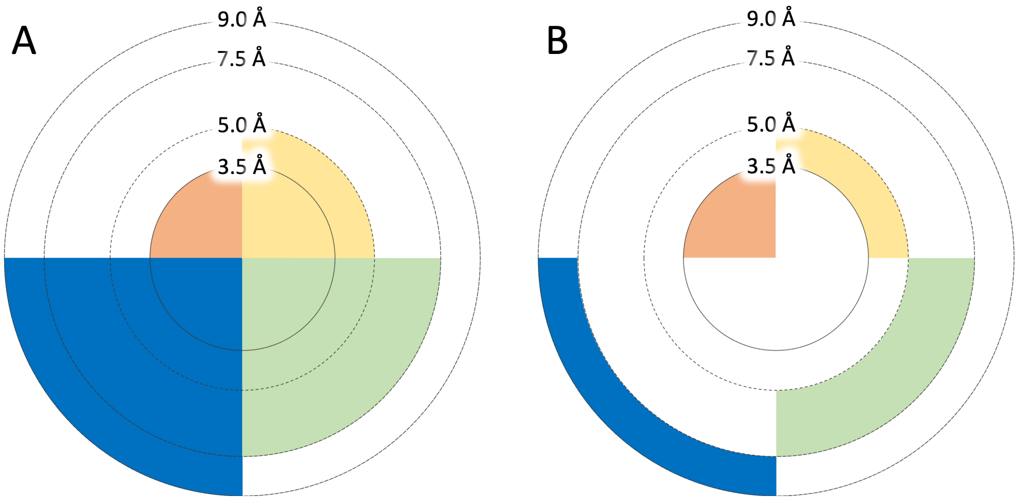

**Supplemental Figure S3: Feature similarity.** (A) Distribution of features used for training. The four groups of Rosetta terms each include 84 features calculated in one of four ways – the mean or average of residues within four shells or spheres – for a total of 294 unique Rosetta category features. (B) Proportion of sites assigned with a filled coordination geometry, coordination geometry with a vacancy, and irregular coordination geometry. (C-F) The feature with the lowest similarity between enzymatic (green) and non-enzymatic (blue) values for the Lining (C), Pocket (D), Electrostatics (E) and Rosetta (F) categories.

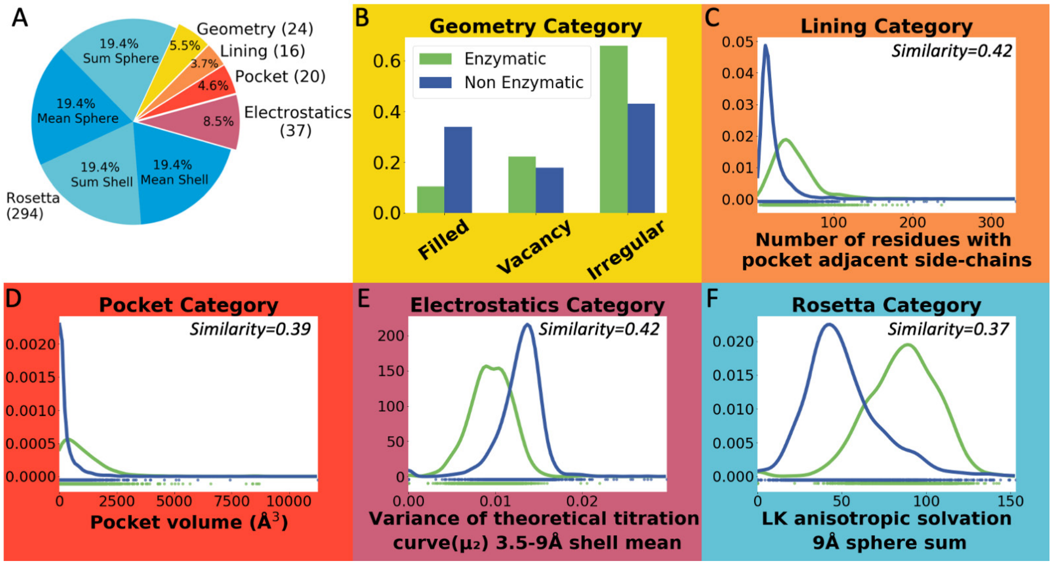

**Supplemental Figure S4: Machine learning experimental design.** The test-set is left for evaluation of the final model while the data-set undergoes nest cross validation (CV). The outer k-fold CV is used for model selections and the inner shuffle split CV is used with grid search to find the optimal machine learning algorithm hyperparameters for each model. Darkest blue is the inner CV training data, medium blue is the outer CV training data, light blue is the outer CV test data and gray is the inner CV test data.

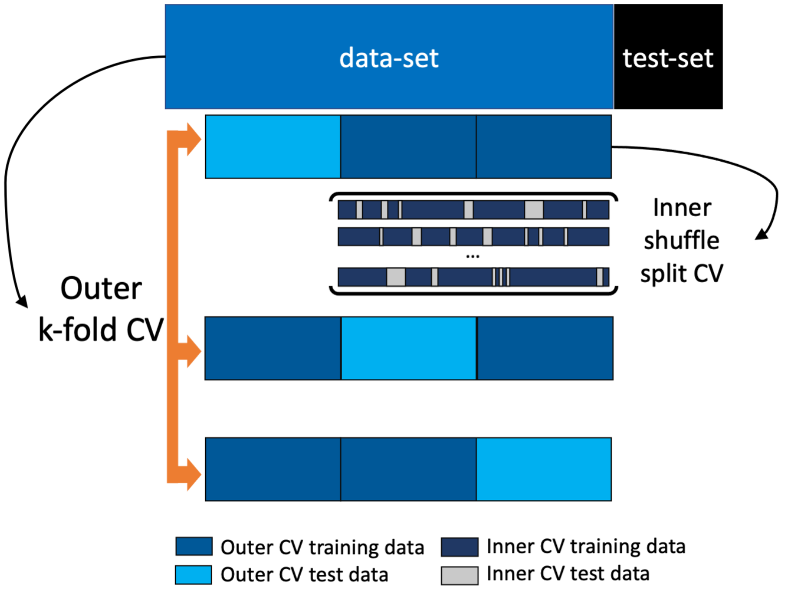


**Supplemental Figure S5: K-Fold deviation.** A kernel density estimates of all models k-fold CV deviation during GridSearch model tuning (left panel) and when held to the single best hyperparameter set (right panel). '---' display standard deviation filters used prior to model selection.

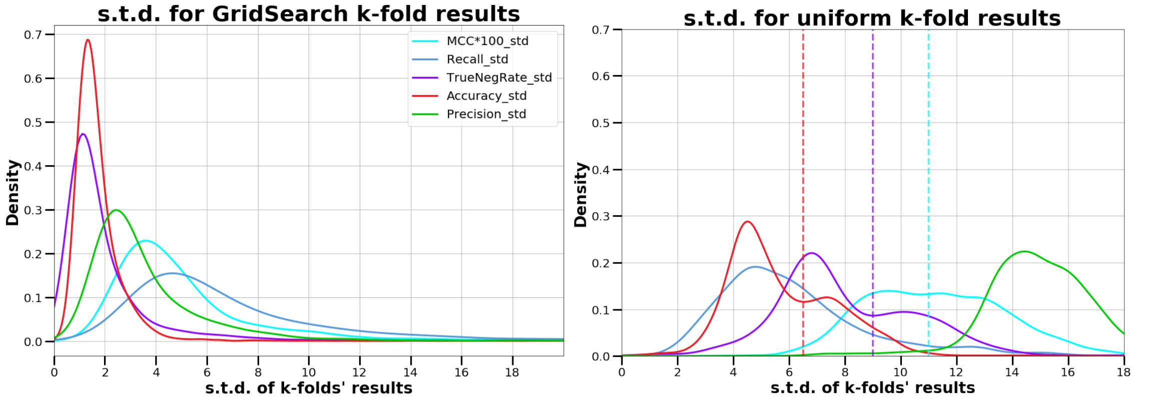

**Supplemental Figure S6: Outer CV performance by optimization metric.** (left panel) Each point represents the results for a specific model after filter (see methods). Points are colored according optimization metric: accuracy (Acc, red), MCC (green), combined MCC and Jaccard index (Multi, light blue), and precision (Prec, purple). Right panel displays zoomed in blue box of left panel.

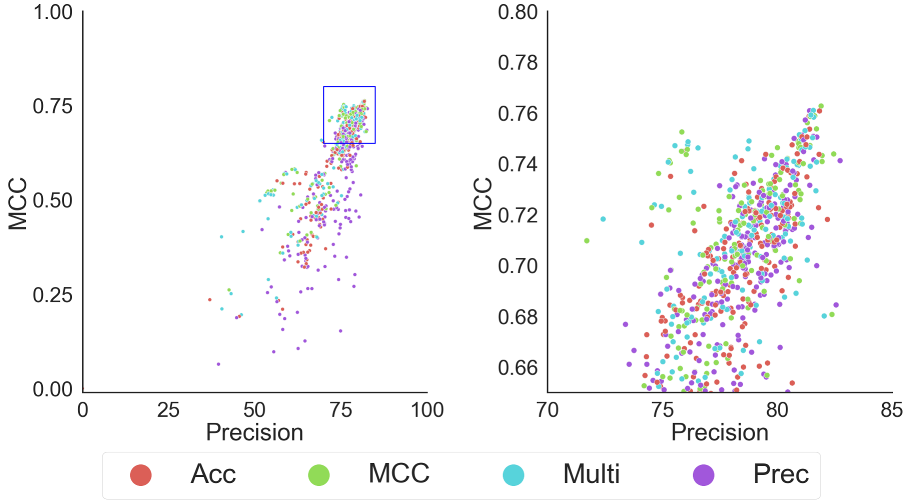


**Supplemental Figure S7: Outer CV performance by different Rosetta category calculation method.** (left panel) Each point represents the results for a specific model after filter which included the Rosetta category in its feature set (see methods). Points are colored according to calculation method: mean sphere (MeanSph, blue), sum shell (SumShell, orange), sum sphere (SumSph, green), and mean shell (MeanShell, red). Right panel displays zoomed in blue box of left panel.

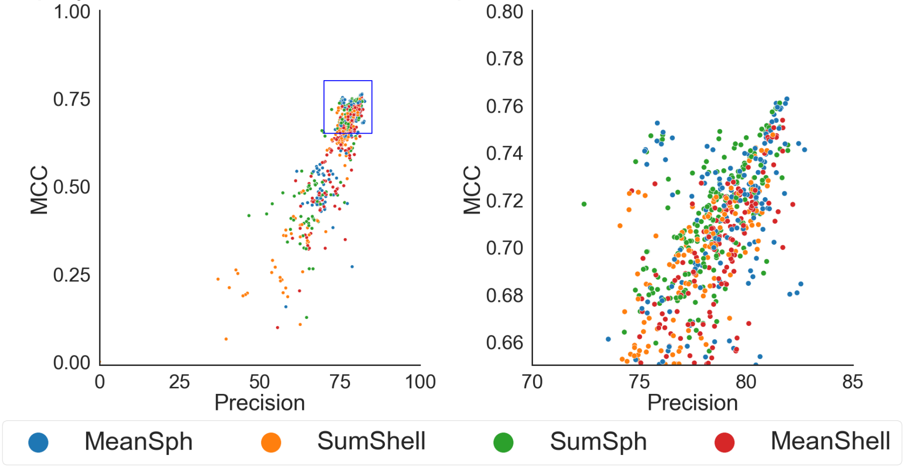

**Supplemental Figure S8: Outer CV performance by feature set.** (left panel) ) Each point represents the results for a specific model after filter. Points are colored according to feature set used by model (Table S2 for feature sets). The size of the dot corresponds to the number of features in that feature set. The right panel zooms in on the blue box in the left panel.  Right panel displays zoomed in blue box of left panel.

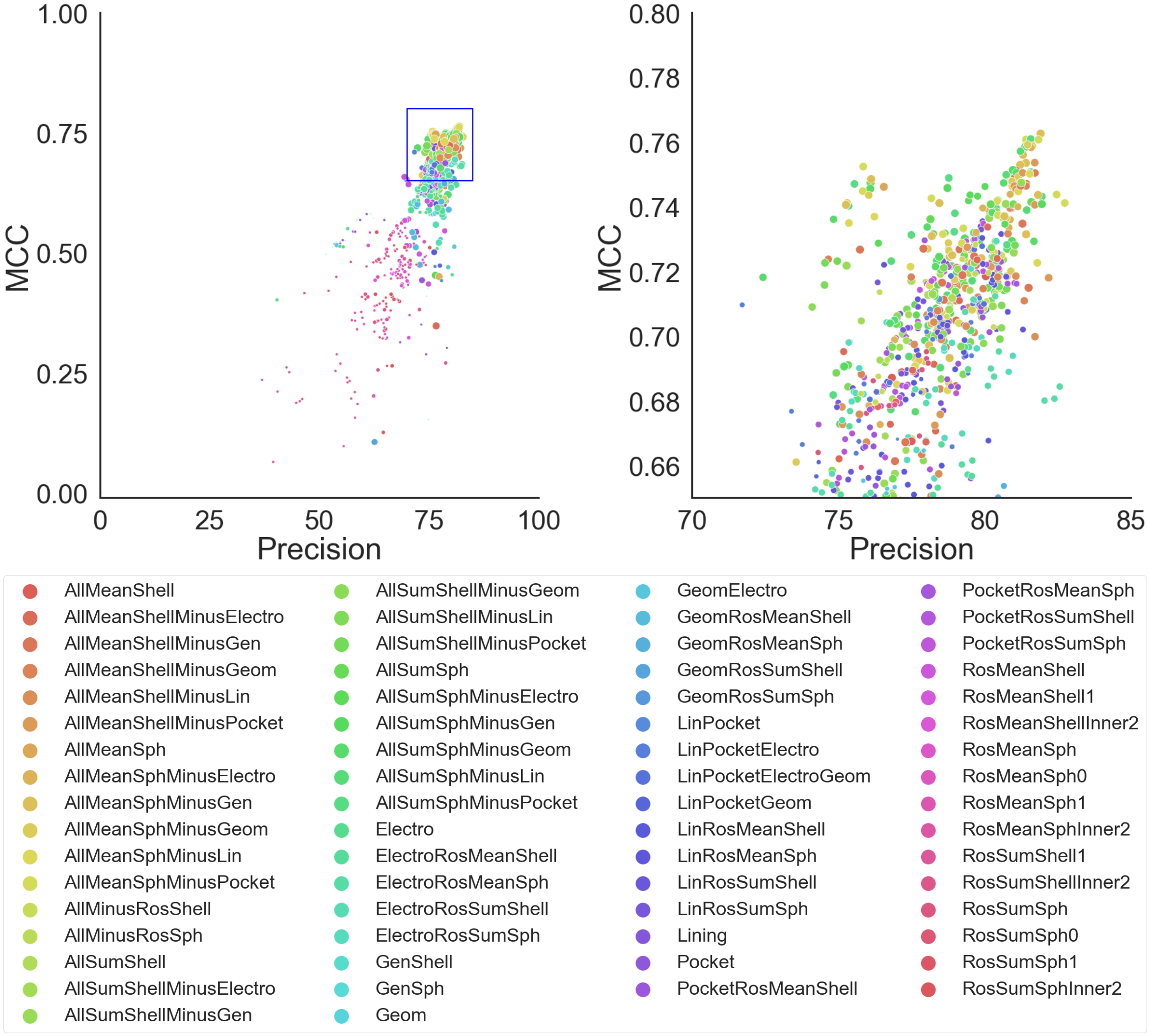

**Supplemental Figure S9: Outer CV performance by algorithm for all models.** (A) Each point represents the results for a specific model. All models are included, even those with high deviation. Points are colored according algorithm used and grouped by classifier type; support vector machines (SVMs) are purples, decision-tree ensemble methods are blues, linear models are reds, discriminant analysis are greens, no grouping for naive Bayes, nearest neighbor, and neural network. Better performing classifiers should be close to the upper right corner. The X denotes our top model (extra trees with AllMeanSph feature set). (B) Zoomed in view of boxed region in A.

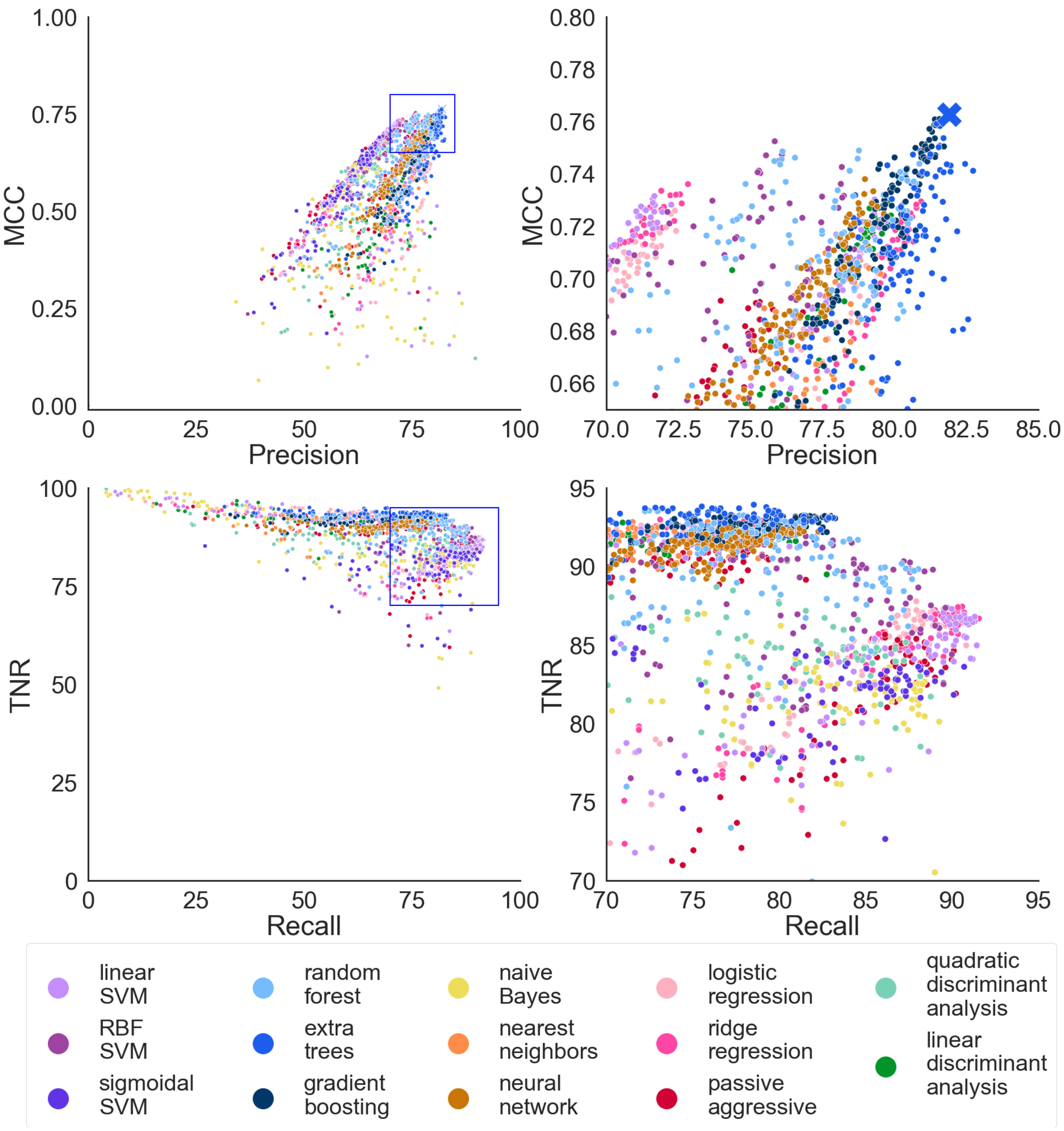

**Supplemental Tables**
Tables ST1, ST2, ST4, ST5, and ST6 are located with more detailed legends in SupplementalTable.xlsx due to size.

**Table ST1: Feature details.** A complete list of features used for machine learning, the category they belong to, and their calculated similarity.

**Table ST2: Feature set details.** All 67 feature set names and the features included in them.

**Table ST3. Performance evaluations for MAHOMES.** The MAHOMES outer CV is the average and standard deviation of the calculated metrics during the second run of our outer, k-fold cross validation (k=7). The other rows show the reported metrics on our hold-out test-set before and after manual check and correction of a few test-set sites.

| Method | Accuracy | Recall | MCC | Precision | True Negative Rate |
|---|---|---|---|---|---|
| MAHOMES outer CV | 90.7% ± 4.4% | 83.1% ± 5.4% | 0.763 ± 0.084 | 81.9% ± 13.8% | 93.1% ± 6.9% |
| MAHOMES test-set | 91.5% | 89.4% | 0.806 | 84.2% | 92.5% |
| MAHOMES test-set corrected* | 94.2% | 90.1% | 0.868 | 92.2% | 96.2% |

**Table ST4: Incorrect predictions.** The test-set sites that were initially incorrectly predicted and our finding from manual inspection.

**Table ST5: Feature importance**. The relative importance output by our extra-trees MAHOMES model

**Table ST6: Prediction results.** The true positives, true, negatives, false positives, and false negatives of the various predictors on their respective T-metal sets.

| Method | True Positives | True Negatives | False Positives | False Negatives |
|---|---|---|---|---|
| **MAHOMES** | 154 | 332 | 13 | 17 |
| **DeepEC** | 152 | 125 | 103 | 16 |
| **DEEPre** | 170 | 184 | 39 | 0 |
| **EFICAz2.5** | 153 | 210 | 20 | 17 |

**Table ST7: Manually annotated M-CSA.** The downloaded M-CSA data with our additional manual annotations. Also includes information for which enzymes and their homologs were removed during the enzymatic labeling process.

**Supplemental Methods**

*Covariate Shift detection*

A covariate shift is when there is a change in the distribution of input data. Developments in methodology to solve protein structures or changes in research funding priority could potentially lead to a covariate shift between our data-set and temporal, holdout test-set. To test for covariate shift, we adjusted our target value to be positive for sites from the test-set and negative for sites from our data-set. We took 300 random positive sites and 300 random negative sites. 80% of this set was used to train a basic random forest classifier using the AllMeanShell feature set, and the remaining 20% was used for evaluation. This process was repeated 100 times, with different random sampling, and resulted in an average MCC value of 0.11, demonstrating that any differences between our test-set and data-set were not substantial enough to be used for differentiating. The process was repeated using catalytic as the target value, which resulted in an average MCC of 0.76, supporting that our methodology could be used to detect significant differences in distribution.

*Feature Calculation PDB Preparation*

Each PDB chain was extracted, including all header information, into a separate PDB file. The PDB was then relaxed [1,2] using the following command line and flags.

```
$ROSETTA_DEV/rosetta_scripts.mpi.linuxgccrelease -s ${pdb_id}.pdb -out:path:all
Outputs/${pdb_id}/ @Relax.flags
```

*Relax.flags:*

```
    -parser:protocol FastRelaxScoreMetals.xml
    -nstruct 50
    -suffix _Relax
    -ignore_zero_occupancy false
    -ignore_unrecognized_res
    -out:file:scorefile Relax.score
    -ex1
    -ex2
    -relax:bb_move false
    -relax:constrain_relax_to_start_coords
    -relax:coord_cst_stdev=0.2
    -beta_nov16
    -overwrite
    -extra_res_fa MO.params NI.params FES.params CUA.params
    -chemical:exclude_patches LowerDNA UpperDNA tyr_diiodinated C_methylamidated
    VirtualBB ShoveBB VirtualDNAPhosphate VirtualNTerm CTermConnect
```

*FastRelaxScoreMetals.xml*

```
<ROSETTASCRIPTS>
        <SCOREFXNS>
                <ScoreFunction name="beta16" weights="beta_nov16" >
                        <Reweight scoretype="metalbinding_constraint" weight="1.0" />
                </ScoreFunction>
        </SCOREFXNS>
        <MOVERS>
```

```
		<SetupMetalsMover name="setup_metals" metals_detection_LJ_multiplier="1.0"
metals_distance_constraint_multiplier="1.2" metals_angle_constraint_multiplier="1.2" />
		<FastRelax name="fast_relax" scorefxn="beta16" />
	</MOVERS>
	<PROTOCOLS>
		<Add mover="setup_metals"/>
		<Add mover="fast_relax"/>
	</PROTOCOLS>
</ROSETTASCRIPTS>
```

Finally, the output structure with the best overall score [3] was kept and scored with no weights as follows:

```
$ROSETTA_DEV/rosetta_scripts.linuxgccrelease -s ${pdb_id}_Relaxed.pdb @Scoring.flags >
StdOutputRelax.txt
```

*Scoring.flags*

```
    -parser:protocol /panfs/pfs.local/work/slusky/MSEAL/RosettaTest/ScoreOnly.xml
    -ignore_zero_occupancy false
    -ignore_unrecognized_res
    -out:file:scorefile Score.score
    -beta_nov16
    -overwrite
    -extra_res_fa /panfs/pfs.local/work/slusky/MSEAL/RosettaTest/CUA.params
    /panfs/pfs.local/work/slusky/MSEAL/RosettaTest/FES.params
    /panfs/pfs.local/work/slusky/MSEAL/RosettaTest/MO.params
    /panfs/pfs.local/work/slusky/MSEAL/RosettaTest/NI.params
```

*ScoreOnly.xml*

```
    <ROSETTASCRIPTS>
        <SCOREFXNS>
                <ScoreFunction name="noweight" weights="EqualWeight"/>
        </SCOREFXNS>
        <FILTERS>
                <BuriedSurfaceArea name="BSA" confidence="0"/>
        </FILTERS>
        <SIMPLE_METRICS>
                <SecondaryStructureMetric name="dssp" dssp_reduced="True"/>
                <PerResidueSasaMetric name="sasa" />
        </SIMPLE_METRICS>
        <MOVERS>
                <ScoreMover name="scoring" scorefxn="noweight"/>
                <RunSimpleMetrics name="run_metrics1" metrics="dssp,sasa" />
        </MOVERS>
        <PROTOCOLS>
                <Add mover="scoring"/>
                <Add mover_name="run_metrics1" />
                <Add filter_name="BSA"/>
        </PROTOCOLS>
        <OUTPUT scorefxn="noweight"/>
    </ROSETTASCRIPTS>
```

#### *Feature Generation*

A number of other programs also generated information from which features were calculated. In all cases, the PDB file is the relaxed structure from Rosetta. The command lines are shown below:

Rosetta pocket_grid [4] for generating the pocket and pocket lining information; rosetta_res is the closest neighboring residue - *$ROSETTA_DEV/pocket_measure.linuxgccrelease -s ${pdb_id}.pdb -central_relax_pdb_num ${rosetta_res} -pocket_num_angles 100 -pocket_dump_pdbs -pocket_filter_by_exemplar 1 -pocket_grid_size 15 -ignore_unrecognized_res*

Bluues [5] for electrostatics information – the pqr file the required input to bluues. We generate it using pdb2pqr.

*#write pqr file*

*/panfs/pfs.local/work/slusky/CommonPrograms/apbs-pdb2pqr/pdb2pqr/pdb2pqr.py --ff=parse --noopt --nodebump ${pdb_id}.pdb ${pdb_id}.pqr*

*#Use bluues to calculate titration curves*

*/panfs/pfs.local/work/slusky/CommonPrograms/bluues/bluues ${pdb_id}.pqr bluues/${pdb_id:0:6}_bluues -pka*

FindGeo [6] for coordination geometry – FindGeo consists of a python wrapper that calls and then parses the output of a pre-compiled program. We extended the python wrapper (see CoordGeomFeatures.py at https://github.com/mwfranklin/CustomModules/) and called FindGeo using the following:

*python $SLUSKY/CommonPrograms/findgeo/findgeo.py -o -p ${pdb_id}.pdb -i ${pdb_id:0:6} -t 3.5*

#### *Machine learning algorithms*

The following algorithms were implemented using scikit learn[7]. All these algorithms are covered in detail as previously published [8].

We chose three support vector machine (SVM) algorithms: linear SVM, radial basis function SVM (RBF SVM), and sigmoidal SVM. SVMs use a set of hyperplanes that separate high dimensional space to create predictions. Inputted feature vectors are mapped into high-dimensional space using a kernel function. Different kernel functions result in different high-dimensional space. We chose SVMs using three different kernel functions: linear, radial basis function, and sigmoidal. Testing different high-dimensional spaces enables different hyperplanes which can perform differently when separating data.

We chose three decision-tree based ensemble methods: random forest, extra trees and gradient boosting. Decision-trees use a series of splits that use a feature to separate the target classes. Splits result in different pathways, or branches, that form a tree-like structure. The ensemble methods combine a group of decision trees to create one prediction. Random forest and extra trees take the average of a set number of independent decision trees. Random forest builds its trees using the best threshold for a feature. Extra trees use random thresholds when splitting. Gradient boosting builds its trees sequentially, allowing new trees to compensate for bias in previously built trees.

We chose three classifiers which fit linear models: logistic regression classifier, ridge classifier, and passive aggressive classifier. Linear models use a set of coefficients, one for each given feature, to generate a prediction. The coefficients are determined by minimizing the residual sum of squares for training data. The logistic regression classifier uses a logistic function to bound the output between 0 and 1, enabling linear models to work for classification. Alternatively, the ridge classifier shrinks the coefficients by adding a size-based penalty. The passive aggressive classifier also uses a penalty but updates after every entry during training.

We chose two models that use discriminant analysis: linear discriminant analysis (LDA) and quadratic discriminant analysis (QDA). Discriminant analysis models probability as the density of a multivariate Gaussian distribution. The predictions for discriminant analysis models follow Bayes' rule.

The remaining algorithms cannot be grouped with each other. Naïve bayes uses the distribution of feature values for each target class to create probabilities which are used for future predictions. Nearest neighbors prediction is based on the class of the most similar feature vector(s) in the training data for its prediction. The neural network implements a multi-layer perceptron, where hidden layers of neurons adjust input data using weights and then transform this data into the output prediction.

**Supplemental Equations**

Equation SE1: Matthews Correlation Coefficient (MCC) calculation [9]. TP is the number of true positives, TN is true negatives, FP is false positives, and FN is false negatives.

$$MCC = \frac{(TP * TN) - (FP * FN)}{\sqrt{(TP + FP)(TP + FN)(TN + FP)(TN + FN)}}$$

**Supplemental Files**

Sites.csv: Final list of sites that were used during the ML process. *SITE_ID* is that site index. *PDB ID* is the crystal structure containing the site. *Chain ID* is the polypeptide identifier for the chain that the site was bound to. *resName1, resName2, resName3,* and *resName4* are the three letter residue codes for the metal ions included in the site, left blank depending on the number of atoms in the site. *seqNum1, seqNum2, seqNum3,* and *seqNum4* are the sequence index in the PDB file for the metal ions included in the site. *Enzyme* is a true for sites identified by our pipeline to be enzymatic and false for those identified as non-enzymatic (the test-set sites found to be mislabeled are not corrected in this file). *Set* is 'data' for sites in the dataset and 'test' for sites in the test-set. *Resolution (Å)* is the crystal structures resolution. *PDB Dep. Date* is the date the structure was deposited in the PDB. *PDB Classification* and *PDB Macromolecular Name* are from the authors of the crystal structure. *EC No* is the downloaded EC number from the RCSB at the time of data collection. *Uniprot Acc* is the accession code(s) in the UniProt KB database for the polypeptide sequence. *Distance Site moved during relax (Å)* is the distance between the site's average location in the original and relaxed structure. *Homolog M-CSA ID* is the entry in the M-CSA database for the sequence homolog with the highest E-value, *M-CSA e_val*, which was further used for enzymatic identification. *Homolog M-CSA aligned TM_len, Homolog M-CSA aligned TM_rmsd, Homolog M-CSA aligned TMscore, M-CSA TM_seqID,* and *Homolog M-CSA aligned catalytic residue distance from site* are the results from aligning the site and its structure with the homolog for the corresponding M-CSA entry.

Test_sequences.txt: The sequence test-set. Each entry starts with a '>' followed by an identifier, which includes uniport accession numbers and one of the SITE_IDs bound to that sequence.

The following Boolean value marks weather it is an enzymatic sequence or not. The next line starts the sequence in FASTA format.
